# Joint identification of sex and sex-linked scaffolds in non-model organisms using low depth sequencing data

**DOI:** 10.1101/2021.03.03.433779

**Authors:** Casia Nursyifa, Anna Brüniche-Olsen, Genis Garcia Erill, Rasmus Heller, Anders Albrechtsen

**Affiliations:** Department of Biology, University of Copenhagen, Denmark

**Keywords:** Autosomes, bioinformatics, resequencing, scaffold-level assembly

## Abstract

Being able to assign sex to individuals and identify autosomal and sex-linked scaffolds are essential in most population genomic analyses. Non-model organisms often have genome assemblies at scaffold level and lack characterization of sex-linked scaffolds. Previous methods to identify sex and sex-linked scaffolds have relied on e.g. sequence similarity between the non-model organism and a closely related species or prior knowledge about the sex of the samples to identify sex-linked scaffolds. In the latter case, the difference in depth of coverage between the autosomes and the sex chromosomes are used. Here we present ‘Sex Assignment Through Coverage’ (SATC), a method to identify sample sex and sex-linked scaffolds from NGS data. The method only requires a scaffold level reference assembly and sampling of both sexes with whole genome sequencing (WGS) data. We use the sequencing depth distribution across scaffolds to jointly identify: i) male and female individuals and ii) sex-linked scaffolds. This is achieved through projecting the scaffold depths into a low-dimensional space using principal component analysis (PCA) and subsequent Gaussian mixture clustering. We demonstrate the applicability of our method using data from five mammal species and a bird species complex. The method is open source and freely available at https://github.com/popgenDK/SATC

## Introduction

The increasing number of non-model organism genome assemblies provide new information on biodiversity and how evolutionary processes have shaped it. An essential part of genome assembly and annotation is the identification of autosomes and sex chromosomes. Eukaryotic species are generally diploid with the majority of their genome represented by autosomes and two sex chromosomes. In mammals the homogametic sex is the female (XX) and the heterogametic sex is the male (XY). This is opposite in birds, where males are homogametic (ZZ) and females heterogametic (ZW). For simplicity we will focus on the XY-system in our method description. Due to their inheritance, sex chromosomes differ from autosomes in several aspects of their population genetics and molecular evolution, e.g. by having a smaller effective population size (*N*_e_) than autosomes (Ellegren 2009) and by having different patterns of population differentiation, especially under incipient or complete speciation (Presgraves 2018). Therefore, it is often preferable to separate them from autosomes in population genetic analyses.

Ideally, whole genome assemblies should be at chromosome level and fully annotated, but due to high cost and challenges associated with complete genome assembly this is often not prioritized for the first generation of a reference genome (Ellegren 2014). Consequently, several approaches to identify sex chromosomes in scaffold level assemblies have been developed (for a review see Palmer *et al.* (2019)). One of them is whole genome synteny alignment (Grabherr *et al.* 2010), where the scaffold-level genome assembly is aligned to chromosome level assembly from a closely related species, and sex-linked scaffolds are identified based on sequence similarity to the reference. There are several obstacles to this approach, most importantly the availability of a chromosome level assembly of a closely related species, but also the accelerated evolution of sex chromosomes in many lineages which causes a high degree of divergence even for closely related species (Charlesworth *et al.* 2018; Irwin 2018; Meisel & Connallon 2013; Presgraves 2018), and computational time (Pennell *et al.* 2018). An alternative is to use genome coverage based methods. These are represented by two groups of methods, e.g. Y-linked and X-linked. Both methods require prior information of the individuals sex, which can be challenging to obtain for species with low sexual-dimorphism, cryptic species, non-invasive sampling etc. Y-linked scaffolds can be identified by mapping sequencing reads from the homogametic sex to a reference genome from the heterogametic sex and the Y-linked scaffolds are identified based on the low depth of coverage (DoC) compared to the autosomes with ½×autosomal DoC in heterogametic sex (Hall *et al.* 2013). X-linked scaffolds can be identified when sequencing reads from both sexes are mapped to a heterogametic reference with the expectation that the homogametic-linked scaffolds will have 1×autosomal DoC in heterogametic sex and ½×autosomal DoC in the homogametic sex. Due to noise in sequencing data the DoC distributions are often overlapping, making it challenging to clearly identify the X and Y scaffolds (Malde *et al.* 2019). The coverage approaches are furthermore highly sensitive to pre-mapping filtering steps and parameters used for the read mapping i.e. repeated regions, average genome-wide DoC etc. (Smeds *et al.* 2015).

Here we present ‘Sex Assignment Through Coverage’ (SATC): a method and software to jointly identify sex-linked scaffolds and determine the sex of each sample mapped to a scaffold level assembly. The method requires sequencing depth information from whole genome resequencing of male and female samples and applies PCA and Gaussian mixture clustering to the DoC among scaffolds to group the dataset into males and females. Hence, the method harnesses the systematic—but noisy—difference in DoC of the sex-linked compared to the autosomal scaffolds. To illustrate how the method works we applied it to five mammal species and a bird species complex with different degrees of DoC, assembly quality (e.g., N50) and sample size. Our method is very fast with computational time being less than a minute for 100 samples. We anticipate that it will be widely useful for inferring individual sex and identification of sex-linked scaffolds for non-model organisms.

## Materials and methods

### Method

The input for SATC are scaffold lengths and mapping statistics (e.g., number of reads mapped to each scaffold for each sample). These are quickly generated with the ‘idxstats’ command in SAMTOOLS (Li *et al.* 2009) from indexed bam files. The method works by i) normalizing the depth of each scaffold within each sample, ii) reducing the dimensionality of the normalized depths using PCA, iii) clustering the samples using Gaussian mixture clustering on the top PCs, and iv) identifying the sample sex and sex-linked scaffolds from the clustering and the DoC. In the first step we calculate for each sample the average depth per scaffold and normalize it by the mean depth of the *M* longest scaffolds in the reference assembly. Suppose we have *s* = 1, 2,…, *S* scaffolds, *n* = 1,…, *N* samples and a matrix RℕSxN containing the number of reads mapped to each scaffold for each sample. We assume that the scaffolds are ordered by length with scaffold 1 being the largest. From this we generate a matrix of normalized mean depth for each scaffold and sample

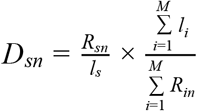

where Rsnis number of mapped reads on scaffold *s,* sample *n* and liis the length of scaffold *i,* i=1, 2,…, M scaffolds. We set M equal to 5. After this normalization, most scaffolds will have a normalized depth close to 1. Following preliminary analyses, we found that filtering out scaffolds < 100kb in length and those with a mean normalized depth outside the range of 0.3 – 2.0 improved the performance of the method, but these thresholds can be set by the user. We then center the matrix by subtracting mean normalized depth from that of each scaffold

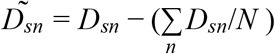

and perform PCA on the **D** matrix. We then identify two clusters of samples that tentatively represent different sex groups (e.g. male and female). For this we use the first two principal components as input for a clustering analysis using Gaussian finite mixture models in *mclust* (Scrucca et al. 2016).

We then identify the sex-linked scaffolds. For this we use the two groups identified by the above clustering and first apply a *t*-test for each scaffold to test for significant differences in mean DoC between groups. We use a Bonferroni corrected *p*-value cut off of 0.05/*S* to identify the sex-linked scaffolds, and scaffolds identified with this test are broadly referred to as sex-linked scaffolds. We then calculate the average depth for each scaffold for both groups. The homogametic (XX/ZZ) and heterogametic (XY/ZW) sexes will show a mean normalized depth ratio of 2:1 between the two groups for scaffolds situated on the X/Z chromosome. Therefore, to identify scaffolds that are situated solely on the X/Z chromosome, we retain sex-linked scaffolds identified with the above method for which the mean difference of normalized DoC between two sex groups is between 0.4 and 0.6. We refer to these scaffolds as X/Z-linked.

For the analyzed dataset in the manuscript the method worked fine without special filtering of the mapped reads or scaffold contents. However, if the user wishes to add additional filters i.e. repeatmasker or removing N’s in the genome assembly, then the input file for the method is easily changed. All analyses were done in R (R Core Team 2019) and are freely available at https://github.com/popgenDK/SATC.

### Application to empirical datasets

To test our method we used the following low to medium DoC datasets mapped to scaffold level assemblies: impala (*Aepyceros melampus*), leopard (*Panthera pardus*), muskox (*Ovibos moschatus*), waterbuck (*Kobus ellipsiprymnus*), gray whale (*Eschrichtius robustus*) and the Darwin’s finches species complex representing 15 species—the mangrove finch (*Camarhynchus heliobates*), the woodpecker finch (*Camarhynchus pallidus*), the small tree finch (*Camarhynchus parvulus*), the medium tree finch (*Camarhynchus pauper*), the large tree finch (*Camarhynchus psittacula*), the gray warbler-finch (*Certhidea fusca*), the green warbler-finch (*Certhidea olivacea*), the Española cactus finch (*Geospiza conirostris*), the sharp-beaked ground finch (*Geospiza difficilis*), the medium ground finch (*Geospiza fortis*), the small ground finch (*Geospiza fuliginosa*), the large ground finch (*Geospiza magnirostris*), the common cactus finch (Geospiza scandens), the Cocos finch (*Pinaroloxias inornate*), and the vegetarian finch (*Platyspiza crassirostris*). We also included two related tanager species; the black-faced grassquit (*Tiaris bicolor*) and the lesser Antillean bullfinch (*Loxigilla noctis*).

The sequencing data from the six different taxa were made available from different studies and were therefore filtered and pre-processed in different ways (see Supplementary Information text S1). Hence, they allow us to assess whether our method is broadly applicable across a range of data treatment regimes, representing the variety of pipelines used in practice for non-model sequencing analysis.

### Validation of sexing and sex-linked scaffolds

To evaluate the sensitivity of the sexing we mapped the five mammal species to a closely related, well-annotated chromosome level reference genome that included an annotated X chromosome. We did not include the Y chromosome because it was not available in all reference genomes and is in general much harder to assemble. For the impala we mapped to the goat (ARS1), for leopard to the domestic cat (Felis_catus_9.0), for waterbuck we used the cow (bosTau8), for muskox we used the sheep (Oar_rambouillet_v1.0) and for gray whale we used the blue whale (mBalMus1.pri.v3). Based on this cross-species mapping we calculated the normalized DoC of X-mapping reads for each individual by the DoC of reads mapping to the five largest autosomal chromosomes of the same external reference genome as an external evaluation of the SATC accuracy in sexing the datasets.

To assess the accuracy and sensitivity of the sex-linked scaffold identification we calculated for each species the sum of the length of scaffolds inferred to be sex-linked or X-linked and compared this number to the length of the X chromosome in the close reference used for validation of samples sex assignment.

## Results

We analyzed whole-genome resequencing data from five mammal species and one bird species complex. The sequencing data was cleaned and mapped to the scaffold-level genome assembly from the same species. The datasets varied both in terms of quality of assembly and in terms of sequencing depth which ranged from 3.13–13.76x. As shown in Table 1 the quality of the genome assembly varied between species, ranging from a scaffold N50 of 344kb in impala to 46.8Mb in muskox and contained 2,796–88,935 scaffolds. These assemblies are representative for many low to medium quality draft genomes. Even after removing scaffolds <100kb a high number of scaffolds remained for some species, up to 7717 for the impala (Table S1).

**Table 1.**
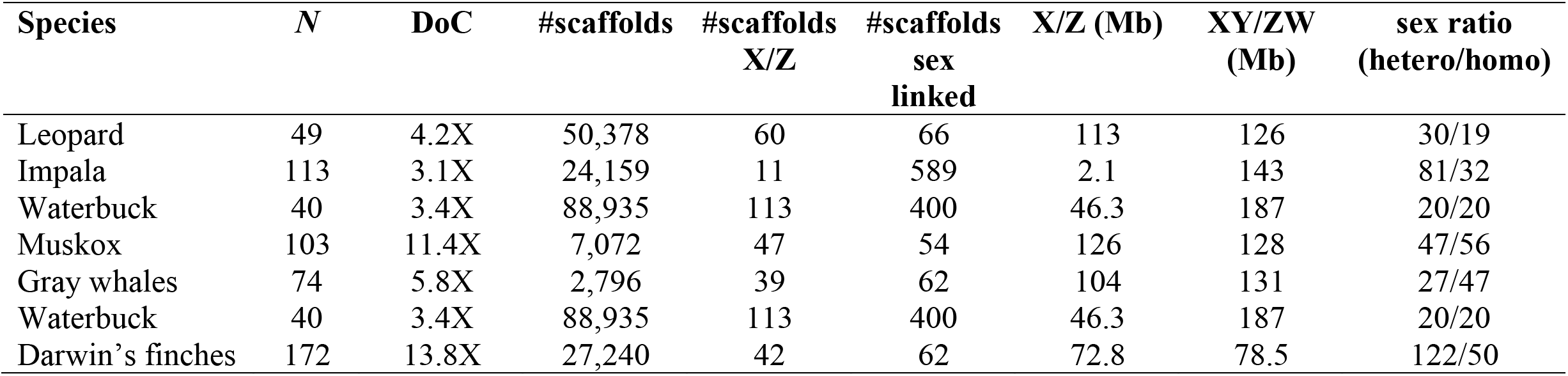
Basic properties of the species analyzed with our SATC method. For each species the number of samples (*N*), depth of coverage (DoC), total number of scaffolds (#scaffolds), number of inferred X or Z scaffolds (#scaffolds X/Z), total numbers of sex-linked scaffold based on t-test (#scaffolds sex linked), length of inferred X/Z scaffolds (X/Z (Mb)), total length of the sex-linked scaffolds (XY/ZW (Mb)), and the sex ratio for the samples. Inferred sex was estimated based on Gaussian mixture clustering from top 2 PCs inferred from closely related species with chromosomal level assembly.

The normalized DoC was very noisy across scaffolds for most species (Fig. S1). However, when we performed a PCA on the DoC matrix we observed for all species a very clear separation into two groups (Fig. 1, left column). For all species except the impala the two distinct groups separated on PC1; for impala this partitioning was on PC2. After applying a Gaussian finite mixture models clustering on the two first axes of the PCA we could clearly group the samples from each species into two groups with characteristic and distinct normalized DoC patterns. We interpret these two groups as the homogametic and heterogametic sex with the homogametic sex being the one with the highest DoC of the sex-linked scaffolds. To validate this we mapped the five mammal species reads to a closely related reference genome containing an annotated X/Z chromosome, and used the normalized DoC of reads mapping to the X chromosome to classify samples as heterogametic or homogametic. This validation showed 100% agreement with the inferred sex using the PCA-based clustering in all cases and across all species (Fig. S3).

**Figure 1:**
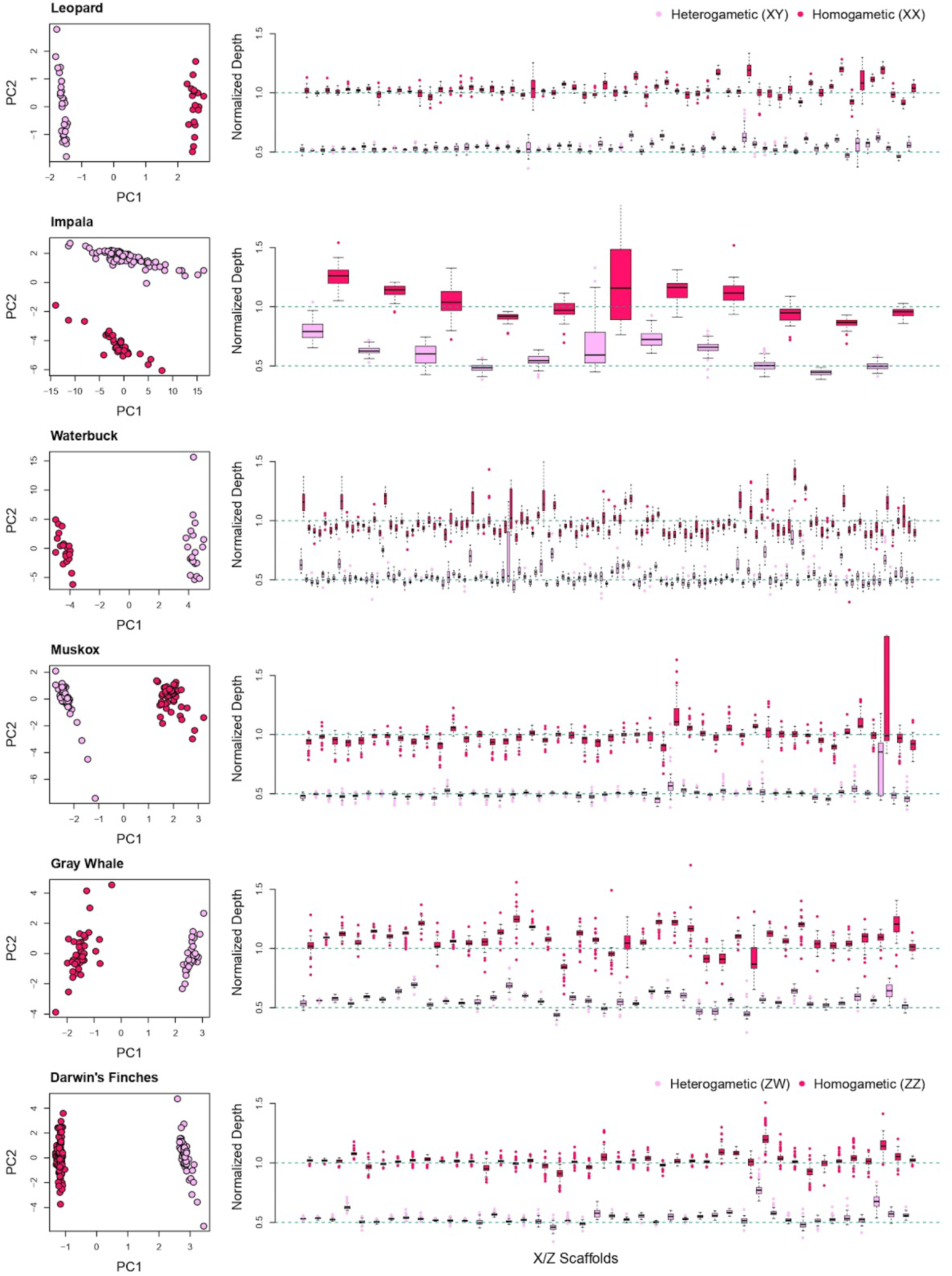
PCA plots with Gaussian mixture clustering and boxplots. Left column: PCA plots of normalized depth across all scaffolds and samples from 5 mammalian species and the 15 species making up Darwin’s finches species complex. Two clusters are inferred homogametic (dark pink) and heterogametic (light pink). Right column: Boxplot of normalized depth from inferred X/Z scaffolds based on mean difference of two sex clusters within range of 0.4 and 0.6. Scaffolds are sorted based on their length (x-axis). Each scaffold is represented by two boxplots from homogametic and heterogametic groups. Expected median values for each group are shown by horizontal green dashed lines of 0.5 (heterogametic) and 1.0 (homogametic).

To identify sex-linked scaffolds we performed for each scaffold a *t*-test for differences in mean DoC in the two sex groups identified above. This identified 54-589 sex-linked scaffolds across the six species, with by far the highest number found in the two most challenging data sets impala and waterbuck (Table 1 and Fig. S2). These sex-linked scaffold might not be exclusively from a sex chromosome and we therefore define them loosely as sex-linked. We furthermore identify X/Z-linked scaffolds by retaining only those sex-linked scaffolds that had a difference of 0.4–0.6 between mean normalized DoC in the two sex groups. This yielded between 11 and 113 X/Z-linked scaffolds in each species (Table 1 and Fig. 1 right column). The total length of sex-linked scaffolds was 126–187 Mb across the five mammals and 79 Mb in the bird species complex, which is close to the length of the assembled X/Z chromosomes for close relatives of each species (Table 1). Many of the sex-linked scaffold that we do not identify as X/Z-linked are scaffolds with DoC that are correlated with sex, but do not adhere to the 0.5:1 ratio. Some of these scaffolds show large deviations from the DoC of other chromosomes while some show an expected Y/W ratio of 0.5:0 (Fig. S2, muskox and Darwin’s finches). Most of the species only had a small number of scaffolds that were sex-linked but not X/Z-linked, however the two species with the most challenging data, impala and waterbuck, showed a much larger difference between the cumulative length of sex-linked and X-linked scaffolds (Table 1). For the impala, the cumulative X-linked scaffold length was just 1.4% of the sex-linked scaffold length, and for the waterbuck this was 24.8%, whereas the corresponding numbers were 79.4-98.4% for the remaining species (Table 1). We therefore conclude that our sex-linked scaffold identification method likely identifies the majority of the X/Z-linked scaffolds while our stricter ratio-based criterion will not identify most of the X/Z-linked scaffolds when reference genome quality and/or resequencing DoC is low.

## Discussion

The increasing amount of whole genome resequencing data present new avenues for population genomic analyses. Herein we add to the analytical toolset by introducing SATC, a method for joint individual sex assignment and identification of sex-linked scaffolds in large datasets. Our method is automated, computationally light, robust to pre-mapping filtering and has a high accuracy. We anticipate that is will be useful particularly for non-model organisms where information on sex of individuals and a chromosome level assembly is often lacking.

The benefits of identifying sex-linked scaffolds when carrying out population genetic studies are many. First, sex chromosomal sites may be desirable for specific analyses, such as association (Lee *et al.* 2017; Luciano *et al.* 2019; Zuo *et al.* 2013), gene expression (Grath & Parsch 2016), or any evolutionary genetic studies on X/Y or Z/W chromosomes (Gottipati *et al.* 2011). Second, if sex-linked scaffolds are not flagged and treated separately they can bias analyses such as demographic history (Li & Durbin 2011), genome scans or genome-wide values of summary statistics, including *F* _ST_ (Lambert *et al.* 2010), genetic diversity (Hammer *et al.* 2010) and allele frequency distribution (Clayton 2008). Third, analyzing males and females separately can elucidate patterns of sex-biased dispersal (Bidon *et al.* 2014) or unequal contributions to offspring diversity (Pérez-González *et al.* 2014).

We show here that PCA on normalized scaffold DoC is a robust approach to identify individual sex for a range of data situations, including having only low-depth resequencing data and a low quality draft assembly as reference genome. Having assigned sample sex we can easily reverse the perspective and utilize this information to identify which scaffolds are sex-linked by exploiting the expected 0.5:1 ratio in sex-specific DoC for each sex-linked scaffold. Our SATC approach does not rely on prior knowledge of sample sex or sex linked scaffolds, and is, to our knowledge, the only automated software that works without any external information.

The recommended usage of SATC would be to flag all scaffolds identified as sex-linked and remove them from further analyses that assume autosomal chromosome data. Conversely, if X or Z-linked sites are desired we recommend to include only those that are flagged as X/Z-linked, i.e. approximately follow the expected 0.5:1 ratio in DoC when compared between the two inferred groups of same-sex samples. Note that it is much harder to identify Y/W-linked scaffolds due to the smaller size of these chromosomes and their highly repetitive sequence content, which can distort the DoC. However, some of the sex-linked scaffolds we identify look like they could be Y/W-linked, having approximately 0.5 normalized DoC in males and very low DoC in females (Fig. S2).

We show that highly fragmented genome assemblies can be used in SATC. The two examined species with the lowest reference genome scaffold N50s (impala and waterbuck) showed deviating patterns from the rest by having very noisy scaffold DoC (Fig. S1). The impala had the lowest-quality genome assembly with a scaffold N50 of 344kb, and for this species we found that the grouping of sexes occurred in PC2 rather than PC1 (Fig. 1). Despite this, our clustering approach was still able to assign sample sex with 100% accuracy. In addition, impalas are known to have segregating karyotypic polymorphisms (Pagacova *et al.* 2011), which could potentially influence the depth across scaffolds and exacerbate the noise in DoC. The waterbuck had a higher than expected amount of sex-linked scaffold content with about 40Mb more content than the X chromosome of the cow reference genome. This could again be influenced by karyotypic polymorphisms, which are known to occur both within and between different subspecies of waterbuck (Kingswood *et al.* 1998). Autosome-to-X translocations are known from several species of bovids (Effron *et al.* 1976; Gallagher Jr & Womack 1992; Kumamoto *et al.* 1996), and if such are segregating within our samples they would complicate the depth-based identification of sex-linked scaffolds. We also observed a large difference between the total amount of sex-linked scaffold content identified by the DoC ratio and the *t*-test methods for these two species, whereas this was much smaller for the other species (Table 1), confirming that excessive noise in scaffold DoC can challenge the use of hard thresholds for identifying sex-linked scaffolds.

Our SATC method also works well with heterogeneous datasets. Darwin’s finches encompass around 18 species of passerine birds (Grant & Grant 2020). We analyzed 15 of these species, which diverged during the last 900-150ky but still have some degree of interspecies gene flow (Lamichhaney *et al.* 2015). Despite the heterogeneous data we were able to assign sex and identify sex-linked scaffolds in the medium ground finch reference genome assembly. We extended the Darwin’s finches dataset with two more distantly related (>900ky) tanager species, the black-faced grassquit and lesser Antillean bullfinch (Lamichhaney *et al.* 2015), and show that the PCA clustering method was still able to reliably assign sample sex as well as identify sex-linked scaffolds (Fig S4).

We emphasize that our method works on data without any need for sophisticated filtering. For example, we did not exclude repeat annotated regions or remove regions without mapped reads prior to calculating the DoC in any of the species datasets. It is possible that additional filtering of the data could improve the identification of sex-linked scaffolds, but we focused instead on demonstrating the robustness of the method by showing that it works in challenging data situations. We found that a single set of settings for the different cutoff values— minimum scaffold length, maximum DoC, ratio of male/female scaffold DoC—yielded usable results for all the species analyzed here. However, the SATC software allows the user to modify these settings if needed. We encourage users to try different cutoff settings to assess the sensitivity of the analyses.

## Acknowledgements

ABO was supported by a Carlsberg Foundation Reintegration Fellowship (CF19-0427). RH, GGE and CN were supported by a DFF Sapere Aude research grant (DFF8049-00098B), and RH was furthermore supported by an ERC Starting Grant (No 853442). AA and GGE are supported by the Lundbeck foundation (R215-2015-4174) and AA the Novo Nordisk Foundation (NNF20OC0061343). We thank the PopGen group at University of Copenhagen for helpful comments on previous versions of the manuscript.

## Data accession

The SRA datasets analyzed in this study are available at the European Nucleotide Archive under the BioProject accession codes: impala (XXXXX), leopard (PRJEB41230), muskox (XXXXX), waterbuck (PRJEB28089), gray whale (XXXXX) and Darwin’s Finches (PRJNA263122 and PRJNA301892). The genome assemblies used were downloaded from NCBI for goat (*Capra hircus*, ARS1, GCA_001704415.1), domestic cat (*Felis catus*, Felis_catus_9.0, GCA_000181335.4), cow (*Bos taurus*, bosTau8, GCA_000003055.4), sheep (*Ovis aries*, Oar_rambouillet_v1.0, GCA_002742125.1), blue whale (*Balaenoptera musculus*, mBalMus1.pri.v3, GenBank assembly accession GCA_009873245.2), gray whale (*Eschrichtius robustus*, XXXXX), the medium ground finch (*G. fortis*, GCF_000277835.1_GeoFor_1.0), impala (*Aepyceros melampus,* IMP GCA_006408695.1), leopard (*Panthera pardus*, PanPar1.0, GCA_001857705.1) and waterbuck (*Kobus ellipsiprymnus*, DFW, GCA_006410655.1). The software framework is freely available at https://github.com/popgenDK/SATC.

## Author contributions

The work was conceived by CN and AA. The research design was planned by CN, ABO, GGE, RH and AA. The data was analyzed by CN and GGE. CN, ABO and RH wrote the manuscript with input from GGE and AA. All authors read and approved the final version of the manuscript.

## References

Bidon T, Janke A, Fain SR, et al. (2014) Brown and polar bear Y chromosomes reveal extensive male-biased gene flow within brother lineages. Molecular Biology and Evolution 31, 1353–1363.

Charlesworth B, Campos JL, Jackson BC (2018) Faster‐X evolution: Theory and evidence from Drosophila. Molecular Ecology 27, 3753–3771.

Clayton D (2008) Testing for association on the X chromosome. Biostatistics 9, 593–600.

Effron M, Bogart M, Kumamoto A, Benirschke K (1976) Chromosome studies in the mammalian subfamily Antilopinae. Genetica 46, 419–444.

Ellegren H (2009) The different levels of genetic diversity in sex chromosomes and autosomes. Trends in Genetics 25, 278–284.

Ellegren H (2014) Genome sequencing and population genomics in non-model organisms. Trends in Ecology & Evolution 29, 51–63.

Gallagher Jr D, Womack J (1992) Chromosome conservation in the Bovidae. Journal of Heredity 83, 287–298.

Gottipati S, Arbiza L, Siepel A, Clark AG, Keinan A (2011) Analyses of X-linked and autosomal genetic variation in population-scale whole genome sequencing. Nature Genetics 43, 741.

Grabherr MG, Russell P, Meyer M, et al. (2010) Genome-wide synteny through highly sensitive sequence alignment: Satsuma. Bioinformatics 26, 1145–1151.

Grant PR, Grant BR (2020) How and why species multiply Princeton University Press.

Grath S, Parsch J (2016) Sex-biased gene expression. Annual Review of Genetics 50, 29–44.

Hall AB, Qi Y, Timoshevskiy V, et al. (2013) Six novel Y chromosome genes in Anophelesmosquitoes discovered by independently sequencing males and females. Bmc Genomics 14, 273.

Hammer MF, Woerner AE, Mendez FL, et al. (2010) The ratio of human X chromosome to autosome diversity is positively correlated with genetic distance from genes. Nature Genetics 42, 830.

Irwin DE (2018) Sex chromosomes and speciation in birds and other ZW systems. Molecular Ecology 27, 3831–3851.

Kingswood S, Kumamoto A, Charter S, Aman R, Ryder O (1998) Brief communication. Centric fusion polymorphisms in waterbuck (Kobus ellipsiprymnus). Journal of Heredity 89, 96–100.

Kumamoto A, Charter S, Houck M, Frahm M (1996) Chromosomes of Damaliscus (Artiodactyla, Bovidae): simple and complex centric fusion rearrangements. Chromosome Research 4, 614–621.

Lambert CA, Connelly CF, Madeoy J, et al. (2010) Highly punctuated patterns of population structure on the X chromosome and implications for African evolutionary history. The American Journal of Human Genetics 86, 34–44.

Lamichhaney S, Berglund J, Almen MS, et al. (2015) Evolution of Darwin’s finches and their beaks revealed by genome sequencing. Nature 518, 371–375.

Lee H-S, Oh H, Yang S-K, et al. (2017) X chromosome-wide association study identifies a susceptibility locus for inflammatory bowel disease in Koreans. Journal of Crohn’s and Colitis 11, 820–830.

Li H, Durbin R (2011) Inference of human population history from individual whole-genome sequences. Nature 475, 493–U484.

Li H, Handsaker B, Wysoker A, et al. (2009) The sequence alignment/map format and SAMtools. Bioinformatics 25, 2078–2079.

Luciano M, Davies G, Summers KM, et al. (2019) The influence of X chromosome variants on trait neuroticism. Molecular Psychiatry, 1–9.

Malde K, Skern R, Glover KA (2019) Using sequencing coverage statistics to identify sex chromosomes in minke whales. arXiv preprint arXiv:1902.06654.

Meisel RP, Connallon T (2013) The faster-X effect: integrating theory and data. Trends in Genetics 29, 537–544.

Pagacova E, Cernohorska H, Kubickova S, Vahala J, Rubes J (2011) Centric fusion polymorphism in captive animals of family Bovidae. Conservation Genetics 12, 71–77.

Palmer DH, Rogers TF, Dean R, Wright AE (2019) How to identify sex chromosomes and their turnover. Molecular Ecology 28, 4709–4724.

Pennell MW, Mank JE, Peichel CL (2018) Transitions in sex determination and sex chromosomes across vertebrate species. Molecular Ecology 27, 3950–3963.

Pérez-González J, Costa V, Santos P, et al. (2014) Males and females contribute unequally to offspring genetic diversity in the polygynandrous mating system of wild boar. Plos One 9, e115394.

Presgraves DC (2018) Evaluating genomic signatures of “the large X‐effect” during complex speciation. Molecular Ecology 27, 3822–3830.

Scrucca L, Fop M, Murphy TB, Raftery AE (2016) mclust 5: clustering, classification and density estimation using Gaussian finite mixture models. The R journal 8, 289.

Smeds L, Warmuth V, Bolivar P, et al. (2015) Evolutionary analysis of the female-specific avian W chromosome. Nature Communications 6.

Team RDC (2019) R: A language and environment for statistical computing, Vienna, Austria.

Zuo L, Wang K, Zhang X, et al. (2013) Sex chromosome-wide association analysis suggested male-specific risk genes for alcohol dependence. Psychiatric Genetics 23, 233.

