## Supplementary Information for "Joint identification of sex and sex-linked scaffolds in non-model organisms using low depth sequencing data"

**Supplementary text S1**

The data sets represent species with different degrees of fragmentation of the assembly and different approaches used for the QC and mapping steps. For the gray whales and Darwin's finches the sequencing reads data were quality controlled using the following steps. TRIMGALORE (Krueger 2015) was applied to remove adaptor sequences and trim consecutive stretches of low quality bases from the ends of the reads. FASTQC (Andrews 2017) was used to generate quality summary statistics for the reads. BWA v0.7.15 was used for mapping reads using the 'mem' function using default settings (Li & Durbin 2010). We validated the sam/bam files and marked duplicates with PICARD-TOOLS v2.9.0 (http://picard.sourceforge.net.). SAMTOOLS v1.4 was employed throughout for bam file manipulation (Li *et al.* 2009). GATK v3.8.0 (McKenna *et al.* 2010) was used for local realignment and duplicate reads removal.

For the impala, muskox and leopards we assessed the overall sequenced reads quality with FASTQC and MultiQC (Ewels *et al.* 2016). We used NGMerge (Gaspar 2018) for impala and leopards, and AdapterRemoval v2 (Schubert *et al.* 2016) for muskox, to collapse overlapping read pairs, requiring a minimum alignment length between read pairs of 11 in both cases. Subsequently we mapped both the merged reads and non-merged reads to the corresponding reference genome for each species with bwa mem v0.7.17 with single end and paired end mode, respectively. SAMTOOLS v1.9 markdup option was used to detect duplicate reads, remove duplicates and low quality mapping reads (-F 3852) in both cases, and keeping only properly paired reads (-f 3) for the non-merged paired end reads. Finally, both merged and non-merged reads were combined into a single bam file. For the muskox we also removed all reads with mapping quality below 30.

For the waterbucks we checked for adapter content using FASTQC and verified mate-pair information and removed duplicate read pairs using Picardtools v2.19. We then identified indel regions using GATK v3.4 and used this information to improve mapping through local realignment.

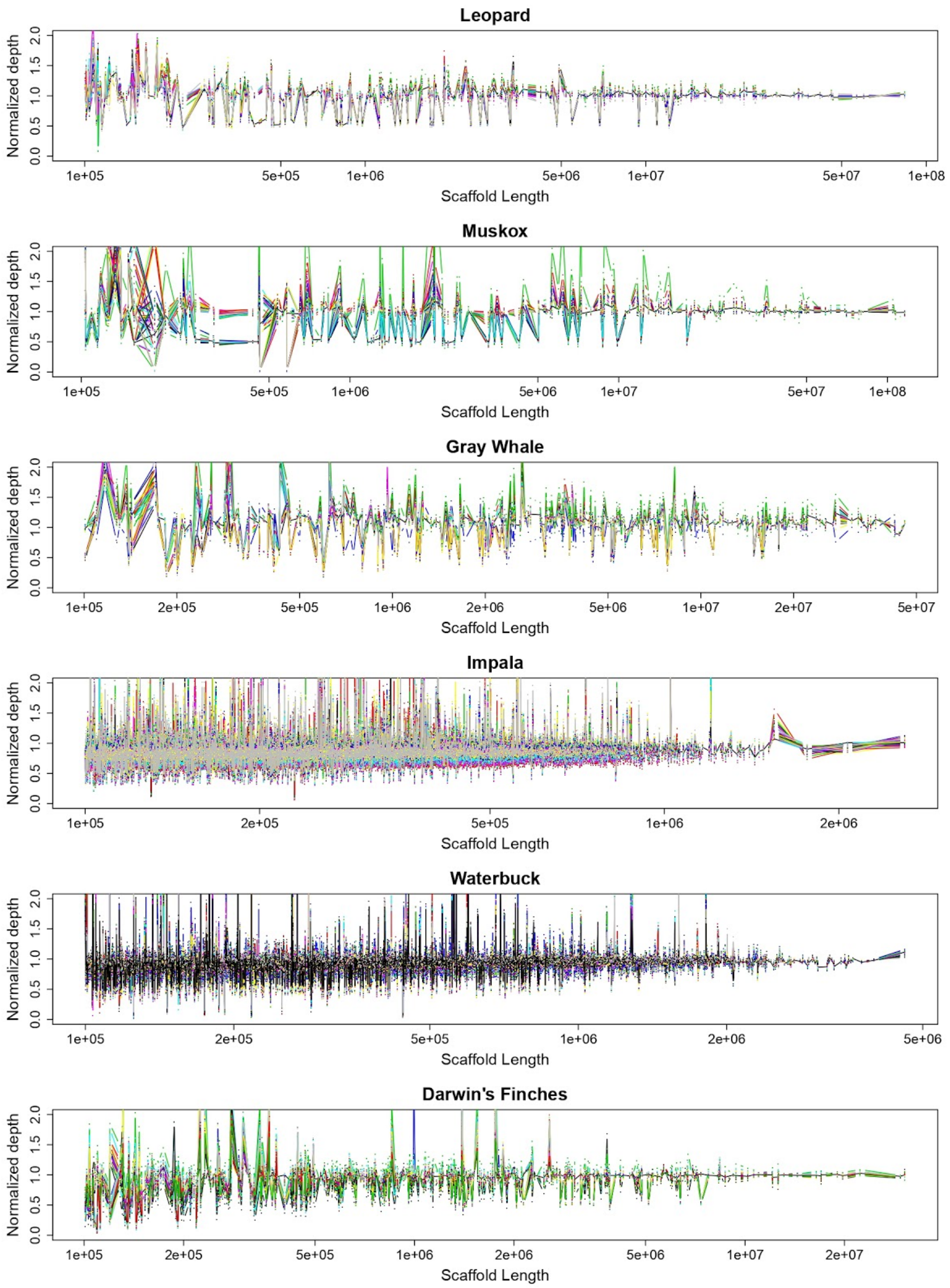


**Figure S1. Normalized depth distribution of >100kb scaffolds from each species.** Scaffolds are sorted according to their length (x-axis). Each line is in a different color and represents normalized depth of each individual across all scaffolds.

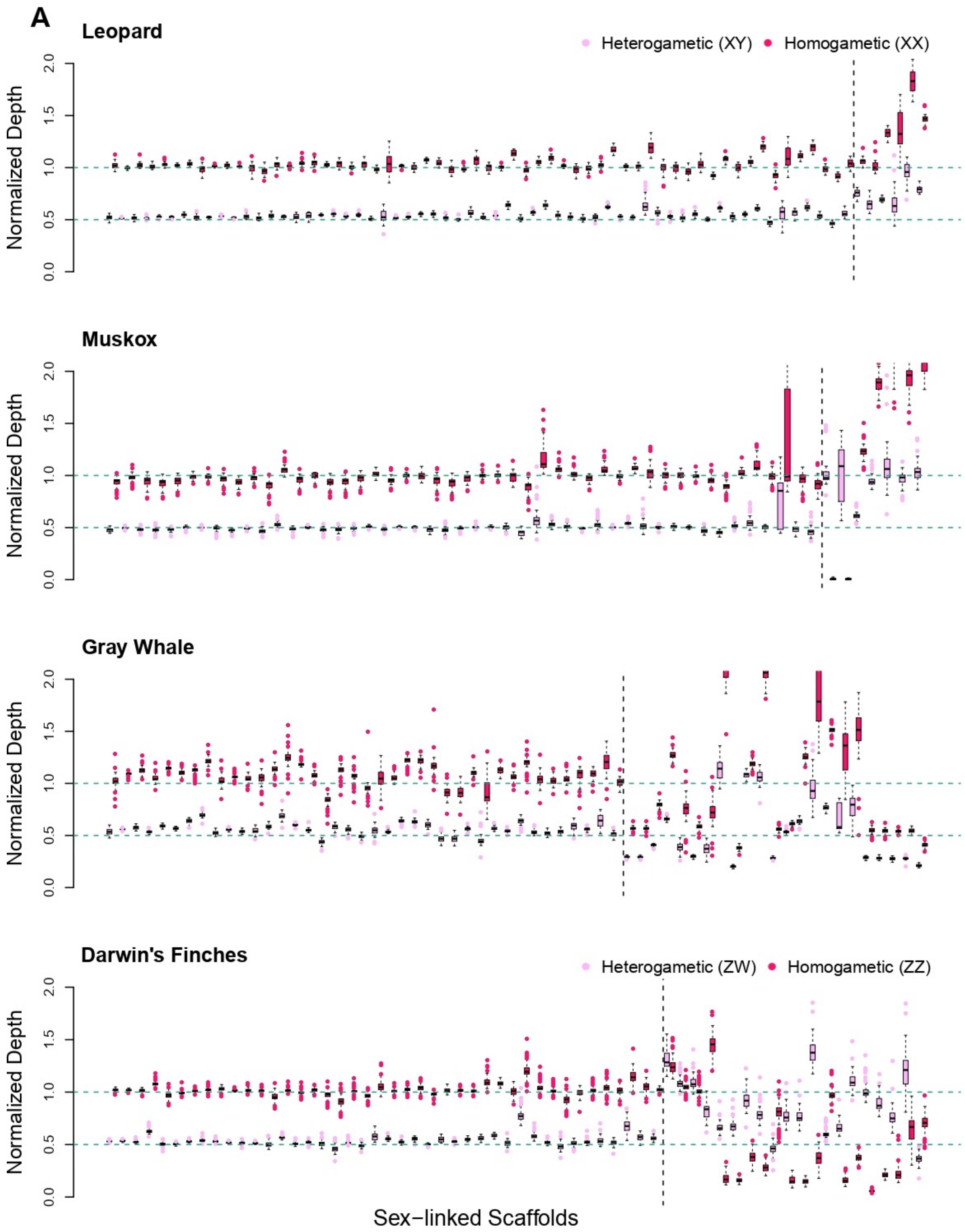
A
Leopard
Heterogametic (XY)
Homogametic (XX)
Normalized Depth
0.0
0.5
1.0
1.5
2.0
Muskox
Gray Whale
Darwin's Finches
Heterogametic (ZW)
Homogametic (ZZ)
Sex−linked Scaffolds

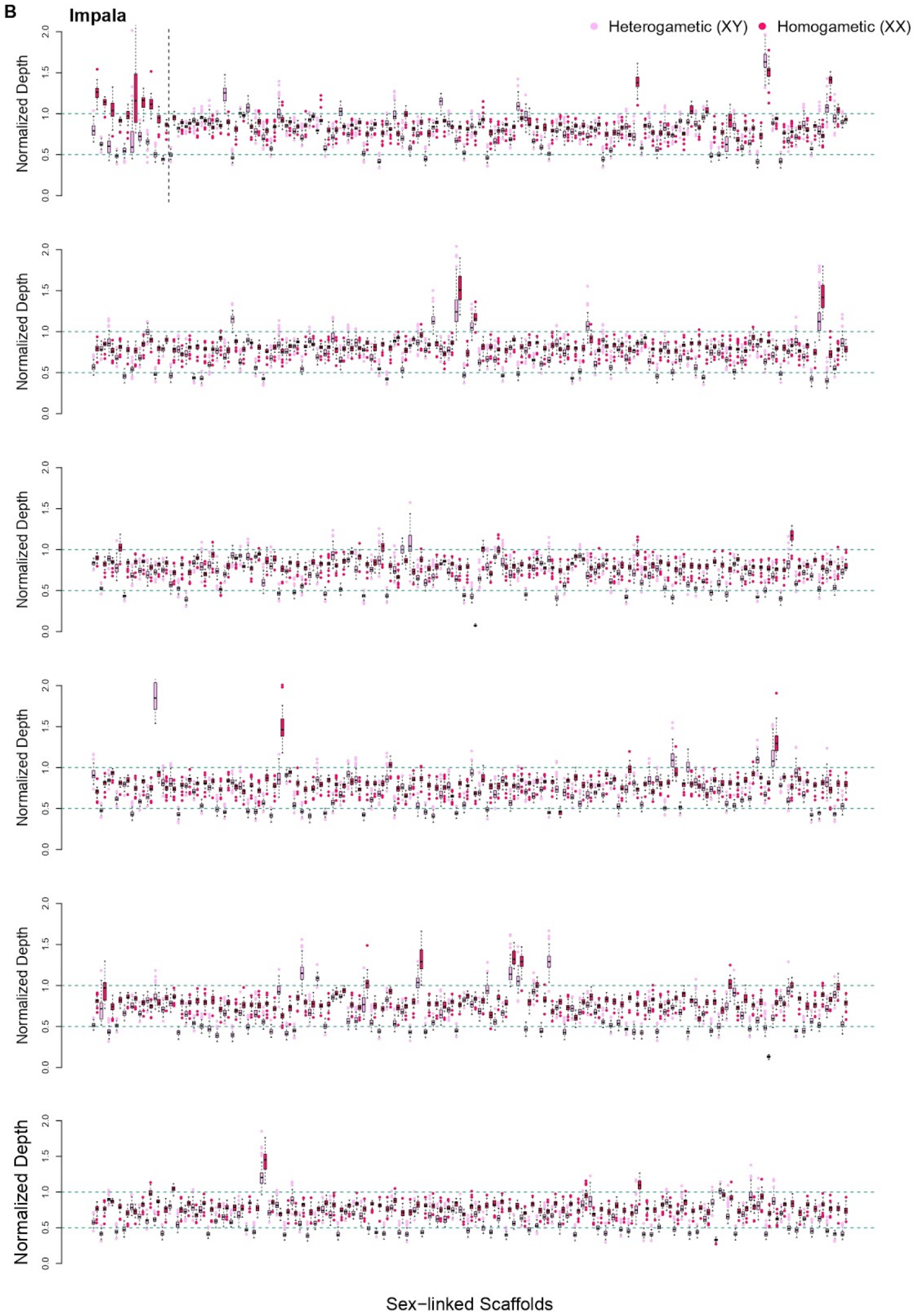
B
Impala
Heterogametic (XY)
Homogametic (XX)
Normalized Depth
0.0
0.5
1.0
1.5
2.0
Sex−linked Scaffolds

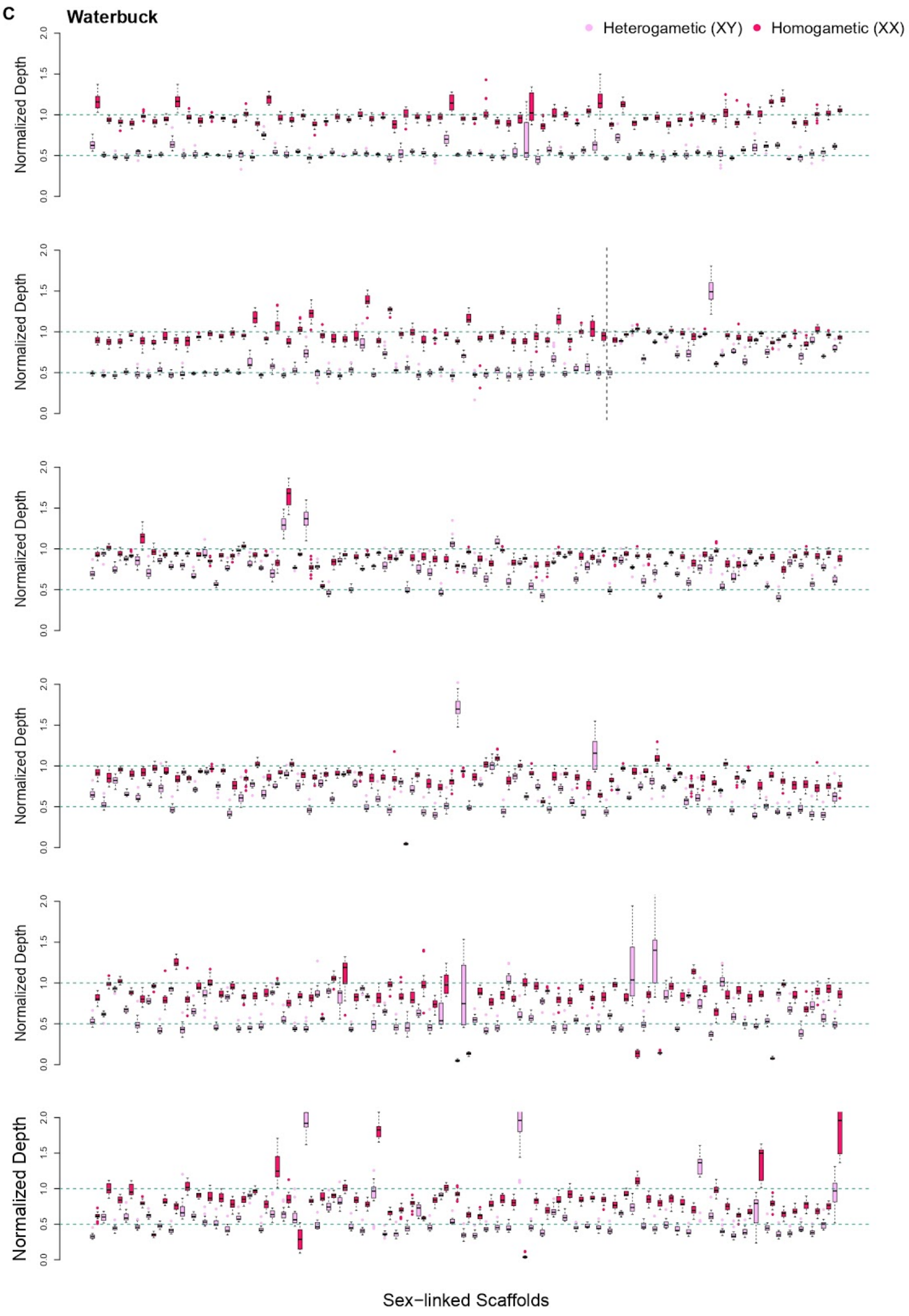
C
Waterbuck
Heterogametic (XY)
Homogametic (XX)
Normalized Depth
0.0
0.5
1.0
1.5
2.0
Sex−linked Scaffolds

**Figure S2. Boxplots of normalized depth of inferred sex-linked scaffolds in (A) leopard, Darwin's finches, muscox and gray whale (B) impala and (C) waterbuck.**

Sex-linked scaffolds are inferred based on mean test (t-test) between two group sex with bonferroni adjusted *p*-value less than 0.05/S where S is number of filtered scaffolds. Scaffolds are sorted based on their length (x-axis). Each scaffold is represented by two boxplots from homogametic and heterogametic groups. Expected median values for each group are shown by horizontal green dashed lines of 0.5 (heterogametic) and 1.0 (homogametic). The vertical dashed line on each species is to separate between scaffolds that pass the t-test and ratio-based threshold (before the line) and scaffolds that only pass t-test (after the line).

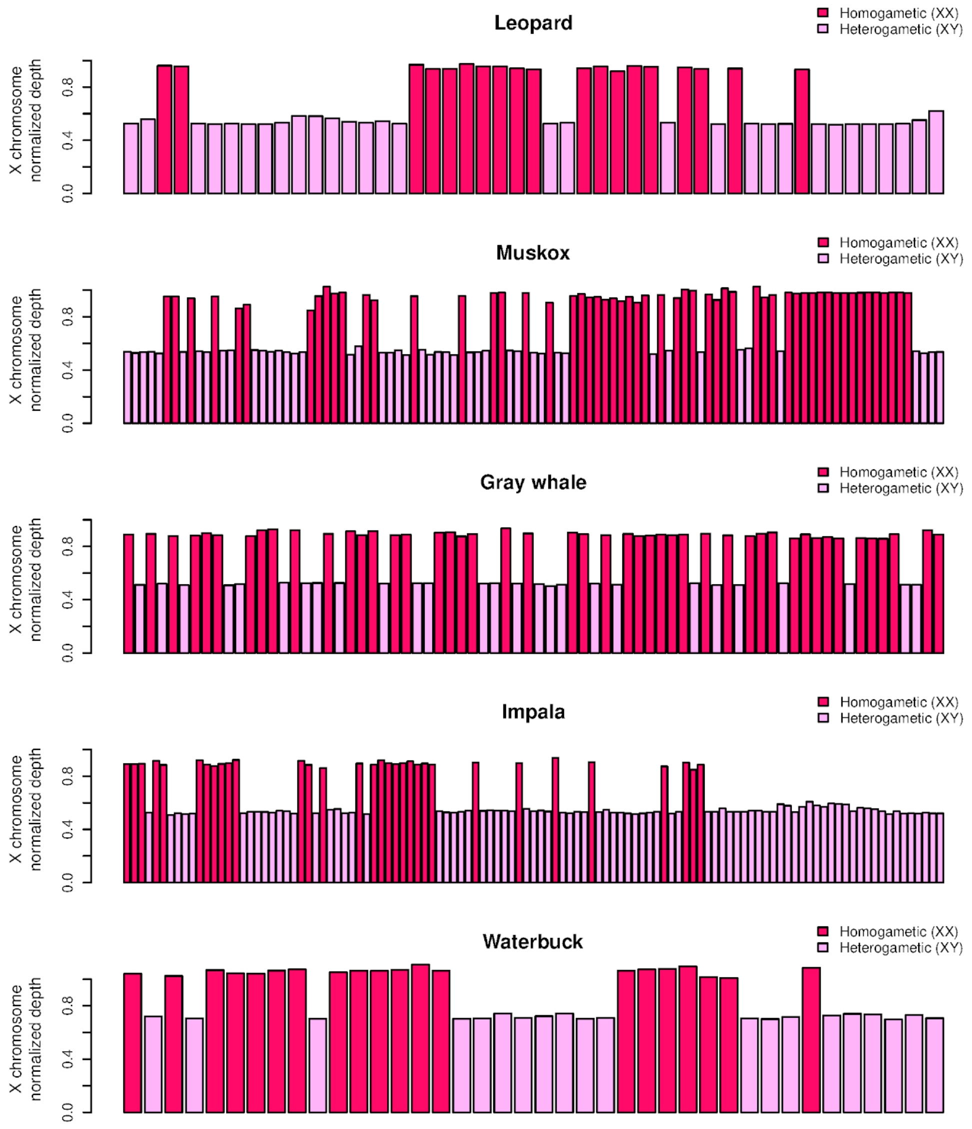


**Figure S3**. **Validation of inferred sex.** Inferred sex from all the samples of each mammalian species based on the normalized depth of reads mapping to the X chromosome of a chromosome level assembly from a closely related species. Samples with X chromosome normalized depth above 0.75 are classified as homogametic and below 0.75 as heterogametic. In all cases samples group in two clearly delimited groups that was consistent with the sex inferred with SATC based on a scaffold-level assembly.

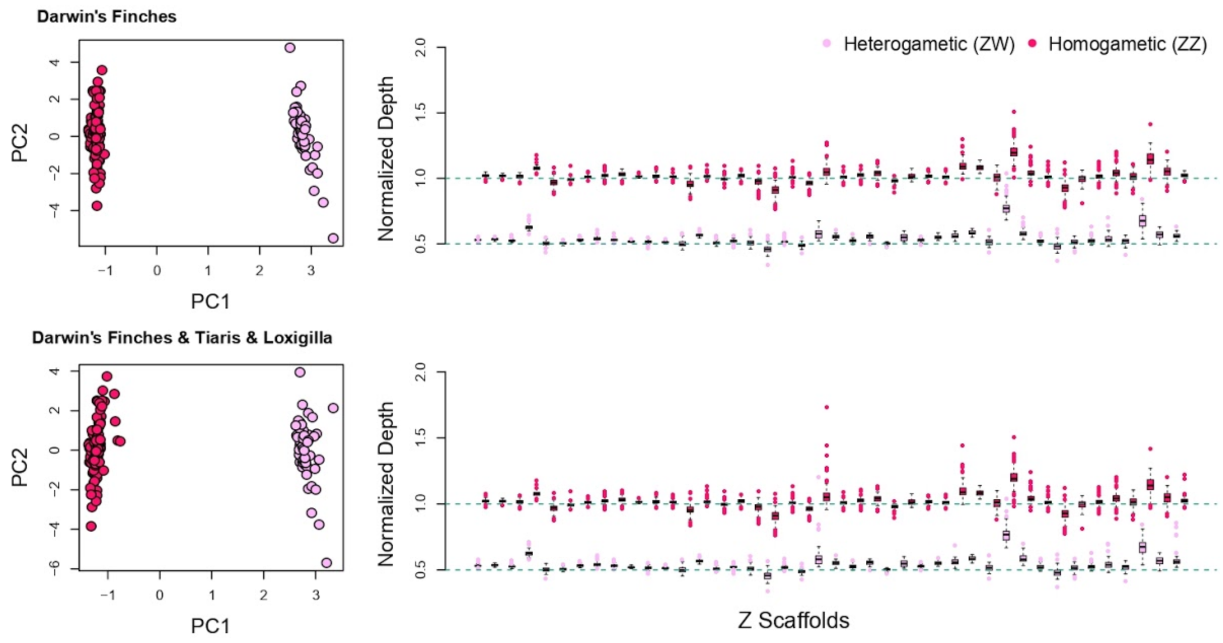


**Figure S4. PCA plots with Gaussian mixture clustering and boxplots for Darwin's finches species complex and two related species.** Plots depicting SATC results for Darwin's species complex and two related tanager species, the black-faced grass-quit (*Tiaris bicolor*) and the lesser Antillean bullfinch (*Loxigilla noctis*). Left column: PCA plots of normalized depth across all scaffolds and samples. Two clusters are inferred homogametic (dark pink) and heterogametic (light pink). Right column: Boxplot of normalized depth from inferred X/Z scaffolds based on mean difference of two sex clusters within range of 0.4 and 0.6. Scaffolds are sorted based on their length (x-axis). Each scaffold is represented by two boxplots from homogametic and heterogametic groups. Expected median values for each group are shown by horizontal green dashed lines of 0.5 (heterogametic) and 1.0 (homogametic).

| Species | Genome assembly length (Gb) | #Scaffolds | #Scaffolds >100K | Scaffolds < 100Kb (Mb) | Length of genome removed |
|---|---|---|---|---|---|
| Leopard | 2.58 | 50,378 | 354 | 33.98 | 1.32% |
| Impala | 2.63 | 24,159 | 7717 | 223.17 | 8.48% |
| Waterbuck | 2.90 | 88,935 | 4608 | 240.00 | 8.29% |
| Muskox | 2.62 | 7,072 | 176 | 37.55 | 1.43% |
| Gray whales | 2.48 | 2,796 | 377 | 32.21 | 1.30% |
| Darwin's finches | 1.07 | 27,240 | 458 | 45.84 | 4.30% |

**Table S1.** Additional statistics on scaffold filtering based on minimum length of 100Kb. We indicated length of genome assembly, total number of scaffolds, total number scaffolds less than 100Kb, total length of scaffolds with length less than 100Kb, and percentage of removed scaffolds.

**References**

Andrews S (2017) *FastQC a quality control tool for high throughput sequence data.* http://www.bioinformatics.babraham.ac.uk/projects/fastqc/

Ewels P, Magnusson M, Lundin S, Käller M (2016) MultiQC: summarize analysis results for multiple tools and samples in a single report. *Bioinformatics* **32**, 3047-3048.

Gaspar JM (2018) NGmerge: merging paired-end reads via novel empirically-derived models of sequencing errors. *Bmc Bioinformatics* **19**, 1-9.

Krueger F (2015) Trim galore. *A wrapper tool around Cutadapt and FastQC to consistently apply quality and adapter trimming to FastQ files* **516**, 517.

Li H, Durbin R (2010) Fast and accurate long-read alignment with Burrows-Wheeler transform. *Bioinformatics* **26**, 589 - 595.

Li H, Handsaker B, Wysoker A*, et al.* (2009) The sequence alignment/map format and SAMtools. *Bioinformatics* **25**, 2078 - 2079.

McKenna A, Hanna M, Banks E*, et al.* (2010) The Genome Analysis Toolkit: a MapReduce framework for analyzing next-generation DNA sequencing data. *Genome Research* **20**, 1297 - 1303.

Schubert M, Lindgreen S, Orlando L (2016) AdapterRemoval v2: rapid adapter trimming, identification, and read merging. *BMC Research Notes* **9**, 88.
